# Stomach serosal arteries distinguish gastric regions of the rat

**DOI:** 10.1101/2021.02.13.431085

**Authors:** Deborah M Jaffey, Logan Chesney, Terry L. Powley

**Affiliations:** Purdue University, 703 Third Street, West Lafayette, IN 47907

**Keywords:** antrum, celiac artery, corpus, duodenum, forestomach

## Abstract

Because the stomach *in situ* has few distinctive surface features and changes shape dramatically with food intake, we have used microCT imaging to (1) characterize the pattern of arteries, potential landmarks, on the stomach wall and (2) evaluate how meal-related shape changes affect the size of the different regions. The stomach receives its blood supply primarily from two pairs of vessels, the gastric and gastroepiploic arteries. Each of the three regions of the stomach is delineated by a distinctive combination of arterial fields: The *corpus*, consistent with its dynamic secretory activity and extensive mucosa, is supplied by extensive arterial trees formed by the left and right gastric arteries. These major arteries course circularly from the lesser towards the greater curvature, distally along both left and right walls of the corpus, and branch rostrally to supply the region. The muscular *antrum* is characterized by smaller arterial branches arising primarily from the right gastroepiploic artery that follows the distal greater curvature and secondarily from small, distally directed arteries supplied by the large vessels of the left and right gastric arteries. The *forestomach*, essentially devoid of mucosal tissue and separated from the corpus by the limiting ridge, is vascularized predominantly by a network of small arteries issued from the left gastroepiploic artery coursing around the proximal greater curvature, as well as from higher order and smaller branches issued by the gastric and celiac arteries. The regions of the stomach empty at different rates, thus changing the dimensions of the organ regions non-linearly.

## INTRODUCTION

Little information is available on the regional patterns of arterial fields located superficially or serosally on the stomach wall. More information is available on the basic pattern of vasculature between the abdominal aorta and the stomach e.g., (Froud, 1959, Greene, 1968, Guth and Leung, 1987, Schnitzlein, 1957, Shukla, 2002, Vdoviakova et al., 2016) and on the microcirculation within the deeper gastric tissues e.g., (Gannon et al., 1982, Guth and Smith, 1975, Holzer et al., 1991, Moskalewski et al., 2002, Peti-Peterdi et al., 1998, Piasecki and Wyatt, 1986, Schnitzlein, 1957), than on the architecture of the vessels running superficially on the stomach wall.

Information about the vessels coursing on surface of the stomach is especially needed because (a) surgeons and physiologists often have to rely on surface features to guide their surgical approach, (b) from the few superficial landmarks that are available, it is difficult to distinguish gastric regions of the stomach *in situ* in the living animal, (c) understanding reproducible patterns might make it practical to implement near arterial infusions of hormonal or pharmacological agents, and (d) nerve bundles such as the abdominal vagus and the sympathetic projections to the gastric vessels throughout the stomach course for a distance near or on the blood vessels as neurovascular bundles, and these nerves could be compromised by vessel ligation or clamping.

More particularly, the rat is considered the most practical and widely used experimental model for surgical research (Lambert, 1965, Martins, 2003, Martins and Neuhaus, 2007, Vdoviakova et al., 2016), and yet the course of serosal blood vessels in the rat stomach (as well as that of other species) has remained almost undescribed. The arterial supply on the stomach wall has conventionally been represented as distributing in schematic, random, and arbitrary generic patterns around the periphery of the organ (see for example, (Greene, 1968, Robert, 1971, Stevens, 1988). For surgeries and physiological experiments, the lack of realistic descriptions and the absence of adequate markers or regional definitions is problematic. Adjustments in size and shape that the stomach undergoes as food is consumed and then macerated for digestion and emptied into the duodenum presumably further complicate assessments of locations for surgical (e.g., bariatric surgeries) access in vivo, attachment of implantable devices, or situating of electrophysiological probes or transducers. Furthermore, few surface landmarks on the organ, other than the borders of the lesser and greater curvature and a hint, from some perspectives, of the deeply situated limiting ridge, are available to guide physiology or surgery of the stomach *in situ*, thus making accurate interventions or placements of electrodes, chemotrodes, strain gauges, or other transducers problematic.

In contrast to the schematic representations, though, we have repeatedly noticed that the major vessels seem to be located in relatively consistent patterns in the rat, offering potential fiducial landmarks that could be used to guide experimental implants or surgeries. In the present experiments, with different degrees of deprivation or food remaining in the stomach, vascular fills of a contrast agent offered a thorough look at the vascular patterns that highlighted certain organizational features of stomach angioarchitecture under different degrees of nutrient load. Large parent vessels reaching the stomach and running on the surface of the organ, in association with the serosa or smooth muscle sheets, routinely filled with contrast agent and were delineated in detail. These vessels appear to distinguish the major gastric regions, irrespective of the distensions and compressions produced by the nutrients within the lumen of the organ. Most notably, the pair of gastric arteries appear to form a regular, particularly recognizable, and lateralized pattern corresponding to the corpus that should be of use in guiding surgical and experimental interventions of the stomach.

## MATERIALS AND METHODS

### Animals

Two- to four-month-old male Sprague-Dawley rats (n =30; RRID:RGD_737903; Envigo, Indianapolis, IN) with an average weight of 277 g (sd = 20 g) were housed individually in shoebox cages with bedding material in an Association for Assessment and Accreditation of Laboratory Animal Care-approved colony room, temperature (22–24 °C)-and-humidity (40–60%) controlled. The room was maintained on a 12:12-hour light–dark schedule. Pelleted chow (2018 Teklad global 18% protein rodent diet; Envigo, Indianapolis, IN) and filtered tap water were provided ad libitum, except during feeding training. All husbandry practices conformed to the NIH *Guide for the Care and Use of Laboratory Animals* (8th edition) and were reviewed and approved by the Purdue University Animal Care and Use Committee. All efforts were made to minimize any suffering as well as the number of animals used.

### Training and feeding

Beginning about one week before the day of each intended experiment, four rats in the desired weight range were trained to consume DietGel Recovery® (ClearH20, Portland, ME). The rats were fasted overnight before being presented with DietGel® at 7am and given 6 hours to consume it, after which the remaining gel was removed and 5g of regular chow was supplied for the rest of the day. After several repetitions the rats naturally consumed DietGel® following an overnight fast. On the morning of the planned experiment, following an overnight fast, a known weight of DietGel® was placed in each animal’s cage; 30 minutes later the bowls containing DietGel® were removed and reweighed to determine the amount of DietGel® consumed by each rat; this time was designated t0. On experimental days, up to three rats consuming at least 6g of DietGel® were then chosen for the experiment; any rat not eating sufficient DietGel® was returned to the general colony population.

### Perfusion and Dissection

Stomachs were stained for microCT analysis in two ways. The first process was designed to stain the entire stomach. An aqueous solution of 0.2% KI and 0.1% I_2_ was made by dissolving KI and I_2_ in ultrapure water. After a delay of 0 to 8 hours from the conclusion of eating, an individual rat was weighed and euthanized with a lethal dose of a combination of ketamine and xylazine ((275 mg/kg ketamine and 27.5 mg/kg xylazine). Once the animal was unresponsive to paw pinch, the abdomen and chest cavity were opened with minimal incisions, and heparin (0.5 ml; 1,000 units/ml) was injected into the heart, followed by transcardial perfusion as follows: 250 ml of 0.01 M sodium phosphate-buffered saline (PBS; pH 7.4; 40°C) was injected with a blunt 18 ga needle threaded via the left ventricle into the aorta (25 ml/min using a peristaltic pump), with blood allowed to drain via a cut in the right atrium and inferior vena cava. During saline perfusion, a 5 Fr or 8 Fr feeding tube was threaded through the esophagus and into the stomach. Similarly, a 5 Fr feeding tube was threaded retrogradely through a small hole cut in the duodenum about 3-4 cm distal to the pylorus and into the stomach via the pylorus. The purpose of the feeding tubes was to ensure that both the pylorus and lower esophageal sphincter remained open to enable subsequent flushing of stomach contents. After exsanguination, the perfusion medium was switched to 500 mL of a fixative solution of 4% solution of paraformaldehyde in PBS. Then, the stomach was removed intact along with approximately 2 cm sections of the esophagus and duodenum, and immersed in the KI/I2 solution on an agitator table. The stomach was held under the surface of the solution and gently manipulated to flush out any Dietgel® and replace the diet with solution, and also to ensure that all air was removed so the organ would not float but be fully immersed for the 24 hour soak period.

The second process was designed to produce a stomach with a stained vasculature. Before perfusion, the Microfil® compound (MV-122; Flow Tech Inc., Carver, MA) and diluent were mixed (about 40 ml total volume, preferred ratio: 3ml MV-122 to 5 ml diluent) in a disposable container and gently shaken or stirred to mix thoroughly but without producing bubbles. The ratio of compound to diluent was chosen empirically to ensure good penetration of the vasculature without the viscosity being so low that the mixture leaked out of the vasculature before curing.

After a delay that was varied systematically from 0 to 8 hours from the conclusion of eating, an individual rat was euthanized and perfused as above. Approximately every minute during perfusion warm saline was administered in and around the abdominal cavity of the rat to minimize cooling. This procedure maximized vasodilation and subsequent penetration of the Microfil® material into the smaller vessels of the forestomach.

After an exsanguination perfusion of around 200 mL saline, the curing agent was added to the Microfil® solution (5% by weight of volume of MV-122 compound (excluding diluent)) and mixed gently but thoroughly for 1 to 2 minutes to achieve homogeneity.

After a total of 250 mL of saline had been perfused, the input to the pump was switched to the Microfil® solution and the pump rate was reduced to 2 mL/min. Warm saline continued to be administered in and around the abdominal cavity of the rat to minimize cooling. Once all the Microfil® solution had been used, the pump was stopped and both the perfusion line and the inferior vena cava (distal to the cut) were clamped to prevent back-leakage.

After about a 70 minute delay to ensure adequate cure of the Microfil® material, the abdomen was opened and the stomach dissected out, gently removing the feeding tubes, and leaving about a centimeter each of esophagus and duodenum attached to the stomach, and also leaving in place the gastroepiploic arteries along the greater curvature of the corpus/antrum and the forestomach. The stomach was then immediately placed in a solution of 4% paraformaldehyde in 0.1M PBS, and left for a period of at least 24 hours.

### Scanning

Before microCT scanning, the stomach (either gastric tissues stained with iodine or vessels filled with Microfil® [or both – see below]) was removed from fix solution and rinsed in saline. To ensure that the stomach was of a reproducible shape and not collapsed during scanning, both duodenum and esophagus were tied off with thread after gently injecting saline into the stomach cavity via the esophagus until it was completely full and contained no air bubbles.

MicroCT scanning was performed with a Quantum GX2 scanner (Perkin Elmer), typically at 90 V /88 A. To enable the stomach to be supported in the preferred orientation without distortion, the stomach was suspended in a viscous solution of Type B gelatin (4-5% in water) in a polypropylene container. The solution was liquid enough that the stomach was fully surrounded by gelatin without pockets of air, but viscous enough that the stomach was fully supported and did not move during scanning. Once scanning was complete, the stomach was thoroughly rinsed in PBS, the tie-off threads were removed, and the stomach was replaced in fixative.

With some Microfil® infused stomachs, following an initial scan to reveal the vasculature alone, the stomach was also lightly stained in iodine solution for about 24 hours, rinsed in PBS for 24 hours to desaturate the iodine stain, and then rescanned to provide an image of both the vasculature and the rest of the stomach.

Scan data was exported as DICOM files and visual images of both the vasculature alone and its combination with the stomach were produced using the open source software platform 3D Slicer (RRID:SCR_005619). In addition, *Vesselucida 360* software (MBF Biosciences, Williston, VT; RRID:SCR_017320) was used to create a 3D reconstruction of the vasculature. Stomach compartment volumes were measured using the *Segment Editor* and *Segment Statistics* modules in 3D Slicer. Images for presentation were generated from both 3D Slicer and from Vesselucida using screen capture and saved as tiff files. For figure generation the tiff files were imported into Photoshop and resized with resampling if needed.

Data associated with this study ((Powley et al., 2020a), (Powley et al., 2020b)) were collected as part of the Stimulating Peripheral Activity to Relieve Conditions (SPARC) project and are available through the SPARC Data Portal (RRID:SCR_017041) under a CC-BY 4.0 license. More detailed experimental protocols are available through Protocols.io (dx.doi.org/10.17504/protocols.io.bafnibme and dx.doi.org/10.17504/protocols.io.95ih84e).

## RESULTS AND DISCUSSION

### RESULTS

It is widely recognized that the basic pattern of blood vessels coursing from the descending aorta to supply the stomach is relatively similar for a variety of species. Between the aorta and the stomach, the pattern in the rat was consistent with the commonly reported arterial distribution. Specifically, the two pairs of major arteries coursing to, and supplying, the stomach are readily recognized: One pair of arteries represent the terminal branches of the celiac artery which divides at the distal esophagus to form the left and right gastric arteries that fork over the lesser curvature close to the lower esophageal sphincter and then branch so as to supply blood to much of the ventral (or left) and dorsal (or right) gastric wall. The right gastric artery often bifurcates from any of several different sites along the more distal branches of the celiac artery.

The second major pair of arteries, the gastroepiploics, issue from higher order branches of the celiac artery and course to perfuse the stomach from the greater curvature. This pair consists of the left gastroepiploic artery running on the rostral or proximal greater curvature and the right gastroepiploic artery coursing around the caudal or distal greater curvature of the antrum. Less conspicuous and apparently less critical are other smaller arteries given off in passing by branches formed from the secondary and tertiary branches of the celiac artery, specifically small arterioles from the splenic artery (to the forestomach) and the gastroduodenal artery (to the antrum).

The overall pattern of the gastric serosal wall angioarachitecture is illustrated in Figure 1. The case illustrated in Figure 1 also appears as an animation (Supplementary Material 1). Notably, the vessels covering the corpus or middle (rostral to caudal) region of the stomach are exceptionally large, presumably capable of delivering considerable blood to the underlying mucosa which is responsible for most of the secretion in the stomach. These lateralized arteries supplying the corpus originate as the left (on the ventral wall) and right (on the dorsal wall) gastric arteries. The gastric arteries travel from the lesser towards the greater curvature along the distal limits of the corpus. As they travel across the stomach, the gastric arteries issue multiple large secondary and higher order branches that distribute out rostrally to supply virtually the entire corpus. The left and right gastric arteries are relatively symmetrical in distribution and comparable in their branching patterns.

**Fig. 1.**
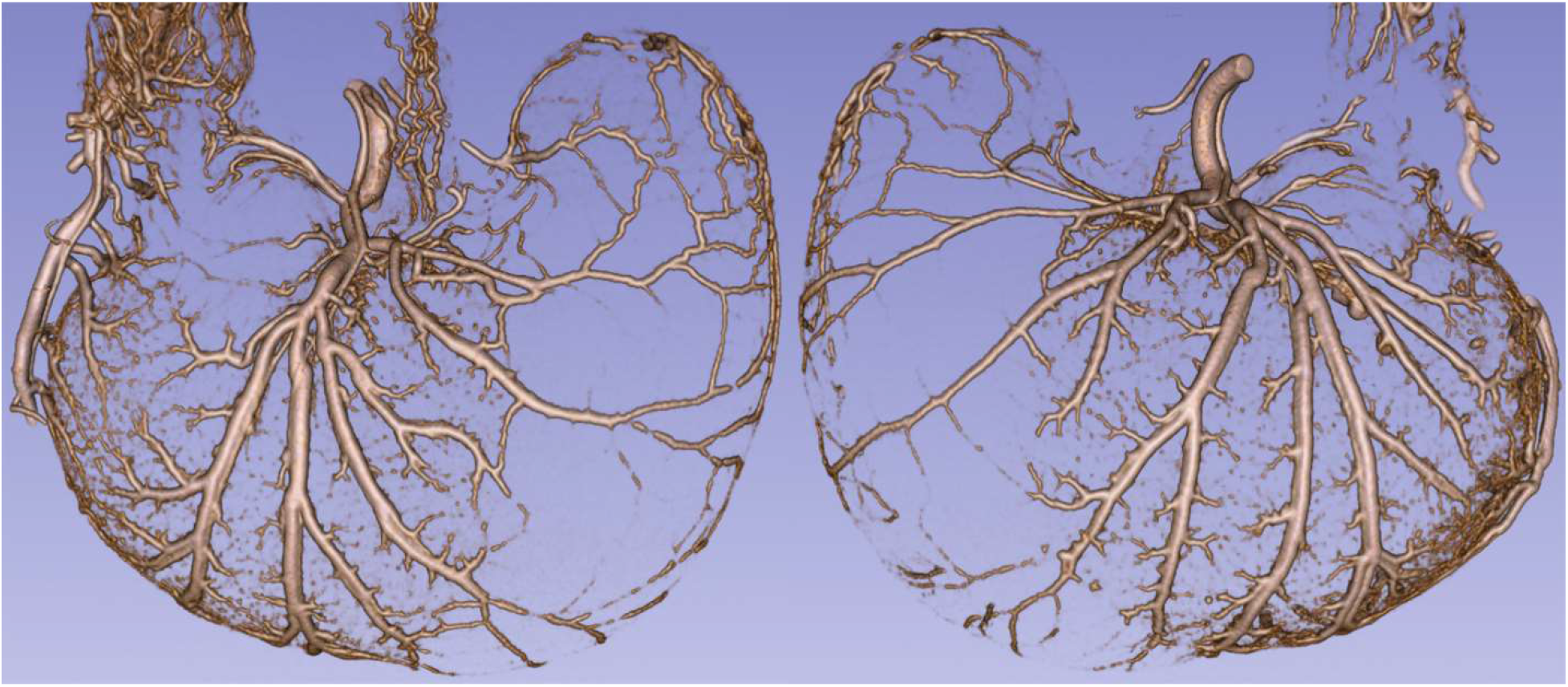
The angioarchitecture profile of the stomach of a recently fed rat delineated with a Microfil^®^ label by microCT imaging (55kV; 144μA). A post-feeding delay of 0.5 hours occurred after a meal of 9.49g DietGel Recovery^®^ was consumed. The profile on left is an exterior microCT view of the ventral stomach wall, and the profile on the right is a comparable exterior view of the dorsal stomach wall of the same animal. The profiles are 3D Slicer reconstructions of microCT scan DICOM files, cropped to include just dorsal or ventral features. The figure was created by screen capture of the 3D Slicer image followed by import into Photoshop where images were resized with resampling as needed. See also Supplementary Material 1 for an animated 3D version.

The gastroepiploic arteries, the other conspicuous pair bifurcating so as to supply blood to the stomach, differ from the gastric arteries in that they are far less symmetrical and are considerably smaller in diameter. The right gastroepiploic artery courses along the distal greater curvature, supplying the antrum and the duodenum with a network of small arteries and arterioles. Along the greater curvature, the right gastroepiploic artery gives off a series of smaller arteries that course towards the lesser curvature or esophagus, supplying the field of the antrum with vascular processes. The bulk of the antral supply of blood seems to arise from the right gastroepiploic artery coursing along the greater curvature, though the antrum also receives some blood supply from the modest numbers of moderately sized higher order arteries that separate from the two gastric arteries.

The left gastroepiploic artery, considerably rostral to the right gastroepiploic artery, courses along the greater curvature of the forestomach, providing smaller, higher-order arteries to the forestomach. Whereas the two gastric arteries are relatively closely matched in size and distributions— basically symmetrical on a sagittal axis—the right and left gastroepiploic arteries distribute at different rostrocaudal levels, making them somewhat asymmetrical on a frontal axis, and are conspicuously different in size. The right gastroepiploic issues medium-sized arteries to both the antrum and (extensively to the) duodenal bulb, whereas the left gastroepiploic artery issues much smaller arteries to the forestomach. The gastric arteries also contribute some vascular distribution to the forestomach, particularly in the “cap” of the forestomach where a conspicuous network of arterioles occurs (see below).

The MBF Biosciences’ software tool *Vesselucida®* was developed to trace the distributions and calibers of blood vessels, and we used it for that purpose. The complementary patterning of the arteries was particularly apparent when we employed *Vesselucida®* to start closer to the celiac artery source and to reconstruct the two conventional pairs of gastric surface arteries. Using *Vesselucida®* we mapped the arterial fields on both the dorsal (left side of top panel) and the ventral (right side of top panel) including the gastric arteries fields (reconstructed in green), the right gastroepiploic artery (reconstructed in red), and the left gastroepiploic artery (reconstructed in purple) separately. Notably too, both of the gastroepiploic arteries are located on the midline and can be seen clearly supplying both the dorsal and ventral surfaces of the stomach. For the animal assessed in Figure 1, Figure 2 (upper panel) displays the *Vesselucida®* patterns of arteries and their relatively distinct perfusion fields, and when the specimen was counterstained with an overnight iodine soak, the shadow of the stomach walls becomes apparent (bottom panel, Figure 2).

**Fig. 2.**
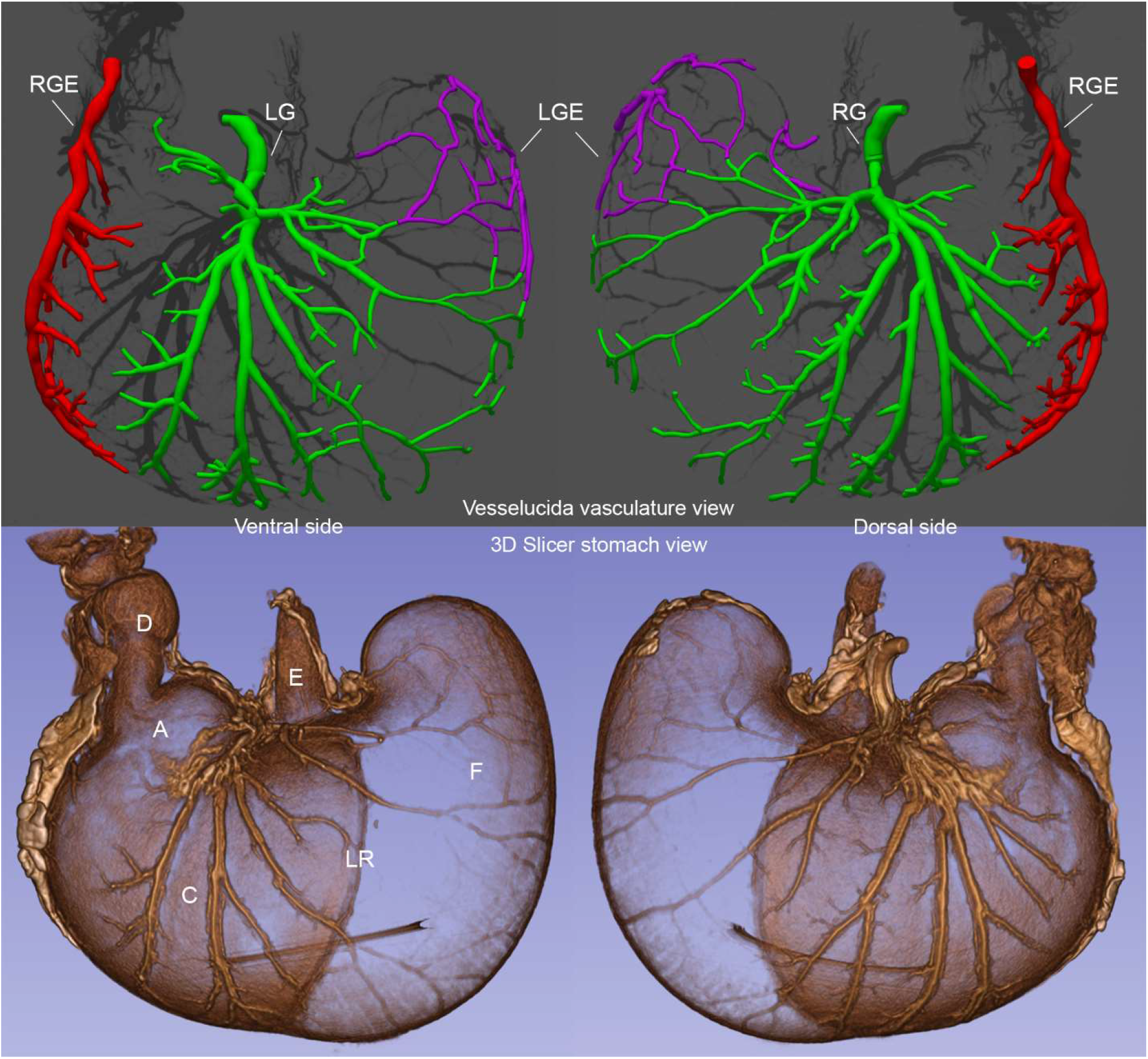
The angioarchitecture profile of the fed rat (same animal as in Figure 1) stomach delineated with a Microfil^®^ label and microCT imaging (55kV; 144μA). A post-feeding delay of 0.5 hours occurred after a meal of 9.49g Dietgel Recovery^®^ was consumed. <u>Upper panel</u>: Profile on left is an exterior microCT view of the ventral stomach arteries mapped and color designated with MBF Biosciences’ *Vesselucid^a®^*, and the profile on the right is a comparable exterior view of the dorsal stomach arteries mapped and color designated with *Vesselucida^®^* of the same animal (3D image cropped to include just dorsal or ventral features). <u>Lower panel</u>: Iodine counterstained profiles of the stomach in the upper panel. Perspectives the same as in the upper panel. The profiles are 3D Slicer reconstructions of microCT scan DICOM files, cropped to include just dorsal or ventral features. The figure was created by screen capture of the 3D Slicer and *Vesselucida^®^* images followed by import into Photoshop where images were resized with resampling as needed. See also Supplementary Material 2 for an animated 3D version. Labels: A: antrum; C: corpus; F: forestomach; LR: limiting ridge; E: esophagus; D: duodenum; LG: left gastric artery; RG: right gastric artery; LGE: left gastroepiploic artery; RGE right gastroepiploic artery.

The stomach stores and partially digests food. Furthermore, the different regions of the stomach, in performing their different functions, have distinctly different thicknesses of wall tissues, muscularity, secretory specializations, and distensibility. An expectation that follows these facts is that, as the stomach empties into the intestines, the different regions of the stomach may empty at different rates. If so, then the stomach might well undergo non-linear adjustments in regional sizes and such changes would presumably complicate surgical and physiological placements on the stomach wall. This effect of differential changes in size can be appreciated by comparing the *Vesselucida®* reconstructions of a stomach perfused when largely emptied of the last meal (6 hr following a 5.94 g meal; volume = 2.7 cm^3^ [close to the minimum observed, c.f. Figure 5]) in Figure 3 with the similar reconstruction in Figure 2 of an animal with considerable food still left in its stomach at perfusion (0.5 hr after a 9.49 g meal; volume = 9.4 cm^3^ [approximately ¾ of maximum volumes observed; c.f. Figure 5]). For example, the heavily muscular antrum and its right gastroepiploic arterial distribution undergoes little size reduction and the secretory corpus and its paired gastric arteries are relatively similar in the comparison of nearly empty stomach in Figure 3 and the relatively full stomach in Figure 2. In contrast, however, the forestomach of the nearly empty stomach in Figure 3 has effectively collapsed producing a dense network of arteries and arterioles in the “cap” region whereas the forestomach and its arteries is relatively distended in the animal in Figures 1 and 2.

**Fig. 3.**
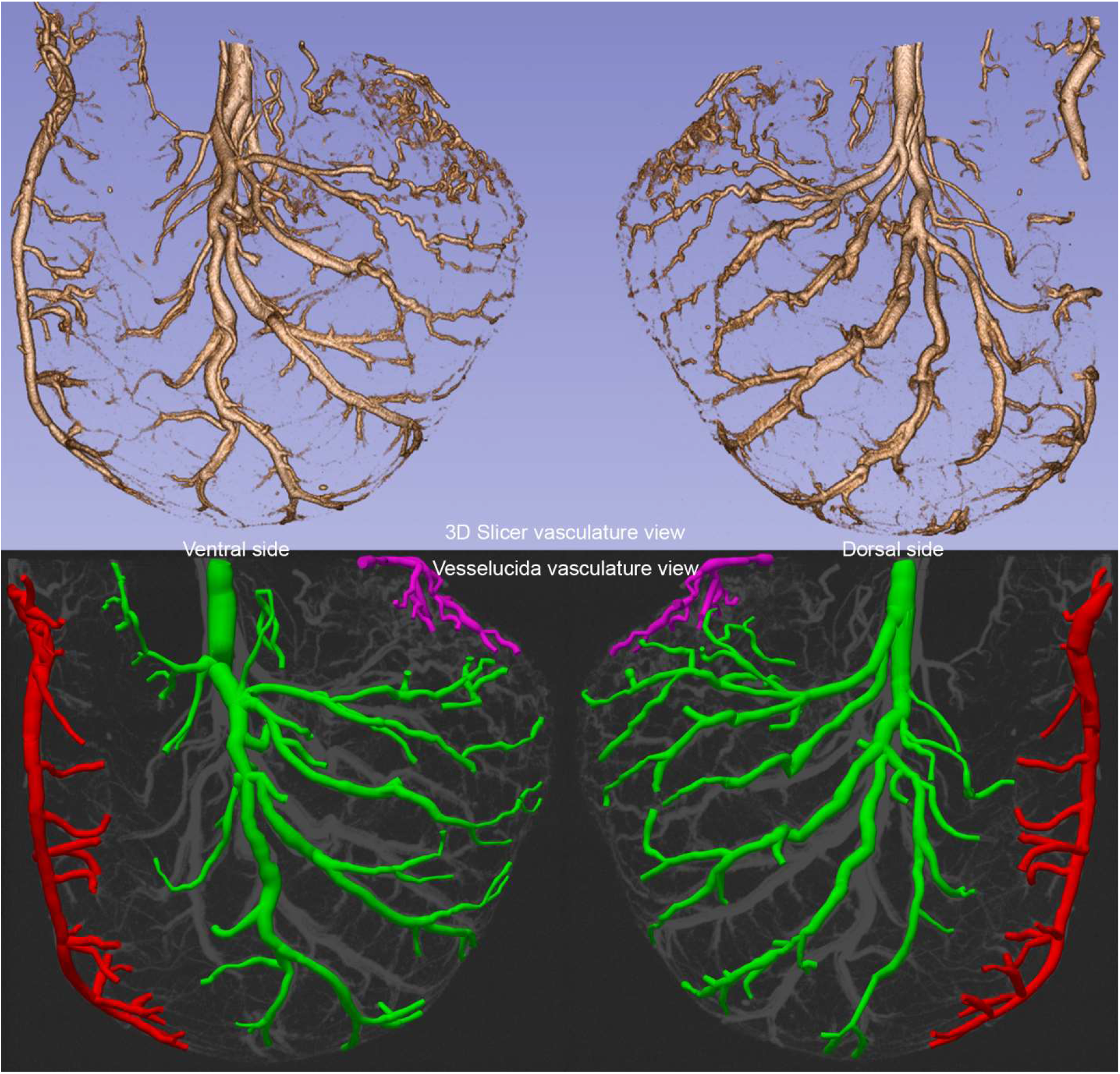
For comparison (with Figures 1 and 2), the microCT angioarchitecture profile of a rat fed (smaller meal of 5.94g of Dietgel Recovery^®^) and given 6 hours for stomach emptying before perfusion and preparation with the Microfil^®^ label; and microCT imaging. <u>Upper panel</u>: Profile on left is an exterior microCT view of the ventral stomach wall arteries, and the profile on the right is a comparable exterior view of the dorsal stomach wall arteries (3D image cropped to include just dorsal or ventral features as appropriate). The profiles are 3D Slicer reconstructions of microCT scan DICOM files. <u>Lower panel</u>: Profile on left is an exterior microCT view of the ventral stomach arteries mapped with MBF Biosciences’ *Vesselucida^®^*, and the profile on the right is a comparable exterior view of the dorsal stomach arteries mapped with *Vesselucida®* of the same animal. The figure was created by screen capture of the 3D Slicer and *Vesselucida®* images followed by import into Photoshop where images were resized with resampling as needed.

We also examined the relationship between the volume of the different gastric regions (presumably related to the amount of food remaining in the stomach) and the size of different meals and the delay times before perfusion, by gauging the size of the overall stomach, as well as the forestomach, and the antrum-corpus (see Figure 4 for illustrative microCT images of iodine stained stomachs of different sizes). Volumes (relative to the outer surface of the stomach since the inner boundary especially in the antrum and corpus was irregular and sometimes indistinct) were measured using *Segment Editor* and *Segment Statistics* modules in 3D Slicer by segmenting the stomach along the limiting ridge. We did not attempt to accurately determine the precise boundary between the corpus and the antrum because (a) this boundary was more indistinct and (b) the forestomach unequivocally emptied faster than the other regions (see Figure 5). Overall, the rate of volume reduction was similar for the forestomach and the entire stomach. The data in Figure 5 correspond to a narrow range of meal size (9-11g) and also include illustrative lines corresponding to a best fit exponential decay model for each set of data (total stomach, forestomach and antrum-corpus, with meal size set to 10g and animal weight set to the study population average of 238.2g). Models were optimized using the entire dataset of 59 stomachs with the *Solver Add-in* within Microsoft Excel (GFG Nonlinear method with multistart option)). The model used in Figure 5 is of the form *v* = *a.W* + *b. M. e*^*c.T + d.w*)^ where v=compartment volume; W=animal weight; M=meal size; T=delay time from meal to perfusion; and *a, b, c*, and *d* are constants. For parameters and statistical assessments of this model and a more complex alternative see Supplementary Material 3. The quantitative model parameters support the idea that emptying of the forestomach is more rapid than emptying of the antrum-corpus (c parameter is more negative for the forestomach than for the antrum-corpus). The minimum or baseline volume of the antrum-corpus is significantly larger than that of the forestomach. This reflects two factors: (1) the sizeable amount of tissue associated with the stomach wall in the corpus in particular; and (2) the consequence of the irregular internal surface of the stomach in the corpus due to the rugae.

**Fig. 4.**
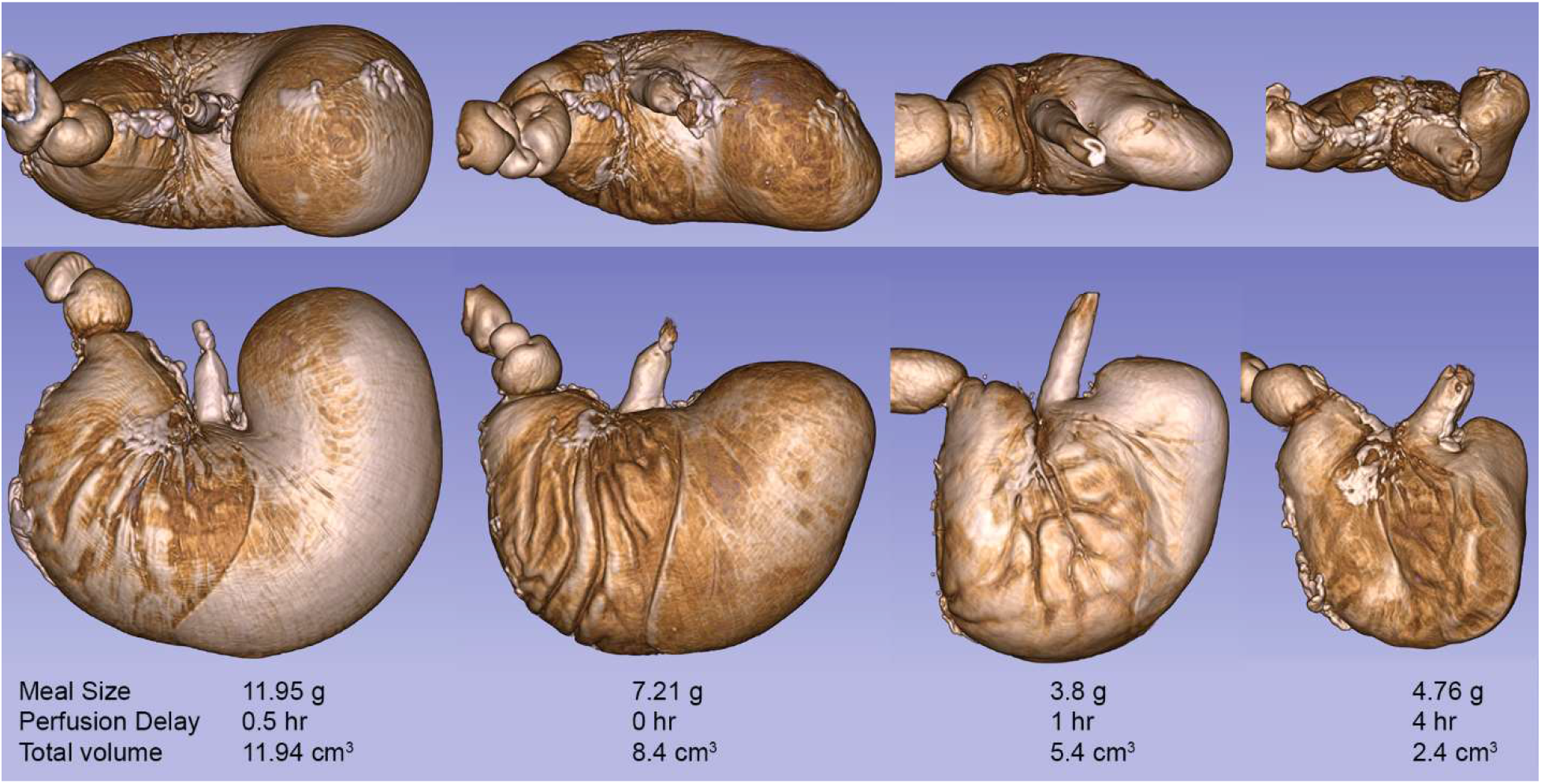
The four sets of views show illustrative microCT images of a variety of iodine stained stomachs (no labeling of the vasculature) at different sizes from full to empty. The images were created in 3D Slicer from DICOM files. Each stomach is labeled to show the meal size, delay time to perfusion and measured total stomach size. The figure was created by screen capture of the 3D Slicer images followed by import into Photoshop where images were resized with resampling as needed.

**Fig. 5.**
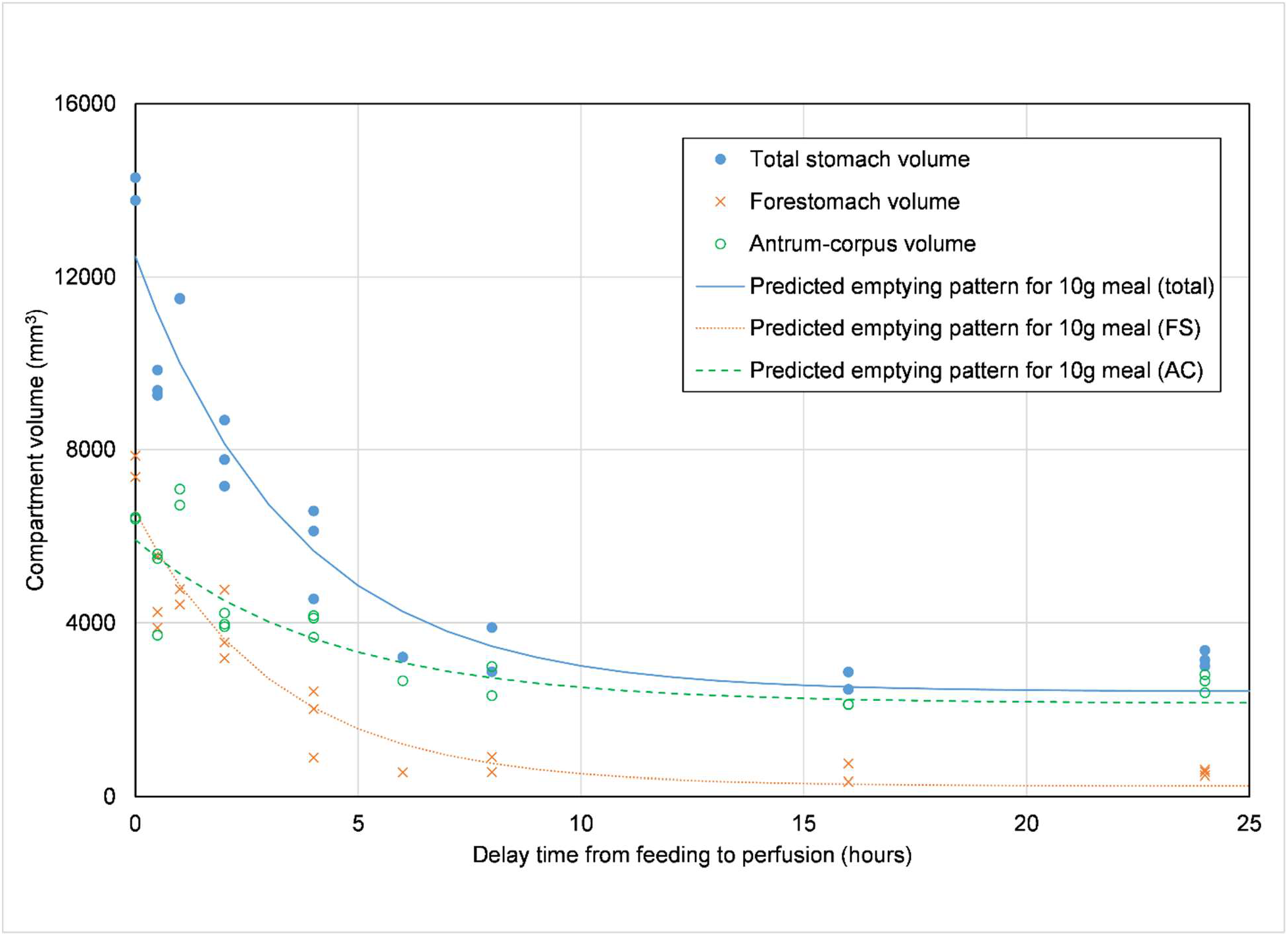
Total stomach volume and compartment volumes (forestomach and antrum/corpus) as a function of the delay time between meal completion and perfusion for meal sizes of 9-11g only, together with illustrative lines corresponding to the fit to an exponential decay model of the form: *v* = *a.W* + *b. M. e*^(*c.T + d.W*)^ where *v* = compartment volume, *W* = animal weight, *M* = meal size, *T=* delay time from meal to perfusion, and *a, b c* and *d* are fit-derived, derived from all the data for each compartment, calculated here for a 10g meal size with W set to the average animal weight for the set of animals. The data shows that the forestomach (which is reflected in the whole stomach emptying pattern) undergoes a dramatic emptying over time (tending towards zero) whereas the antrum-corpus undergoes a much more modest emptying pattern consistent with the observation that food is stored primarily in the forestomach and digested and emptied—but not accumulated—in the antrum-corpus.

A final observation—one we had not anticipated—also deserves mention: We had not anticipated the degree to which small arterioles in the mucosa would apparently respond to a newly ingested meal. Immediately before perfusion, animals were given a half hour to eat plus a subsequent period for emptying. A few of the animals (11) were perfused at t = 0, or immediately after the half hour of access to DietGel Recovery®. Of those animals, some (n =3) had weak or incomplete vascular fills. Of the remaining 8 animals with complete vascular fills 6/8 exhibited a dramatic filling of the fine arterioles and vessels of the mucosa, illustrated in Figure 6, in which the inset panel demonstrates a high level of fill, in contrast to that seen, for example, in Figure 3 where the animal was perfused several hours after eating. The pattern is consistent with the idea that consumption of nutrients after an overnight fast triggers an initial gastric phase of digestion which involves a dramatic secretory response from the mucosal lining of the corpus. In the animals that were perfused anywhere from a half hour to an hour and a half after the meal, the dilation of the mucosal arterioles was already considerably reduced.

**Fig. 6.**
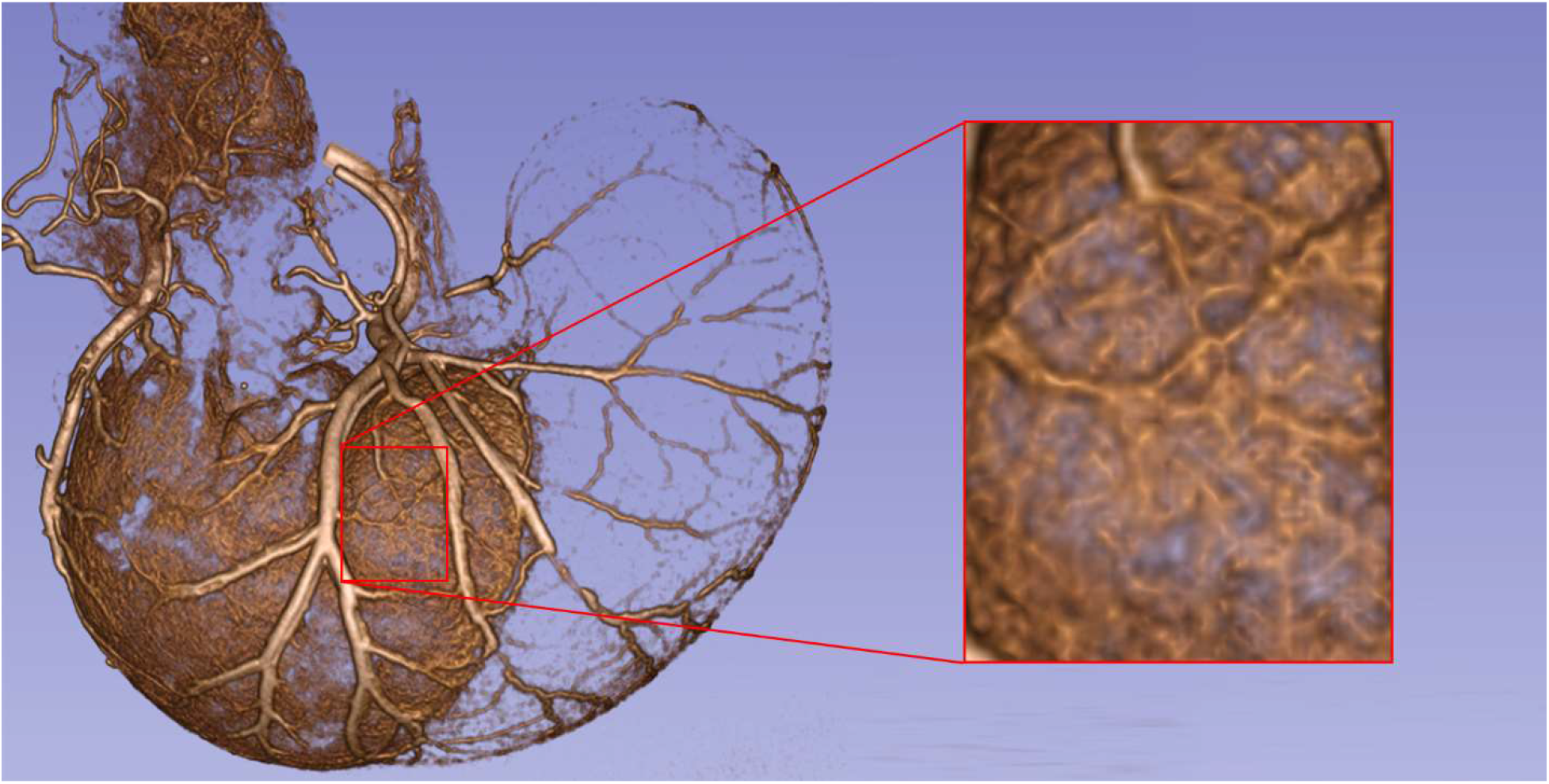
Animal perfused immediately after the 30-min feeding period. 5.3g Gel consumed. Animals perfused immediately after feeding characteristically exhibited labeling or filling of fine arterioles in the gastric mucosa throughout its distribution distinguished by the large vessels of the gastric arteries (see inset). The profile is a 3D Slicer reconstruction of microCT scan DICOM files. The figure was created by screen capture of the 3D Slicer images followed by import into Photoshop where images were resized with resampling as needed.

## DISCUSSION

During rodent surgeries—experimental bariatric analyses, electrophysiology, physiology, nerve isolations, biopsies, etc.—from the relatively undifferentiated appearance of the surface of the stomach, it is often difficult to determine the regions such as the antrum, corpus or forestomach cf. (Robert, 1971). Other than the overall profile of the organ (e.g., the greater and lesser curvature) and, at least in some illuminations, the limiting ridge in the gastric wall, the stomach is hard to divide unequivocally into regions e.g., (Lambert, 1965).

Ironically, conventional morphological sources have largely ignored the arterial architecture observed on the serosal sides of the stomach wall. Some sources ignore the vasculature of the stomach altogether e.g., (Stevens, 1988), others simply illustrate an inaccurate annulus of small vessels entering the stomach at the greater and lesser curvatures e.g., (Robert, 1971), and still others arbitrarily suggest larger vessels on the wall of the stomach that do not appear to be close approximations of the actually architecture e.g., (Greene, 1968). Ironically, while the Greene text provides what is still perhaps the most comprehensive description of the angioarchitecture of the rat (152 total pgs., 177 – 329) “circulatory system”, it only devotes one figure (# 246) and nine lines of text to the arteries and veins connecting the descending aorta and the vena cava, respectively, to the stomach. The Greene volume does not attempt to characterize or name the arteries (or veins) on the surface of the stomach wall.

Furthermore, the lack of distinguishing markers that definitively identify the particular regions of the stomach is further complicated by the fact that the organ, in its role as a storage vessel, varies dramatically in size and regional dimensions as a function of deprivation or feeding (see Figures. 4 and 5). Indeed, we speculate that this variability is partially responsible for apparent variability in the angioarchitecture of the stomach. To address the lack of adequate descriptions of the arterial supply of the stomach and potentially to address complications introduced by the changes in gastric shape associated with differences in gastric fill, we have examined the superficial arterial vessels of the stomach. The results suggest that the major vessels of the two canonical pairs of arteries supplying the stomach with blood might be used as fiducial markers to enhance reliable surgical exposures and placements for stomach interventions.

### Angioarchitecture of Rat Stomach Surface

Under the methodological conditions outlined above, in ongoing experiments in the laboratory, a regular and instructive angioarchitecture of the rat gastric vasculature was observed. Each of the three major divisions of the stomach, namely the antrum, corpus and forestomach, was well delineated by a distinctive and characteristic arterial field. The three fields were not so rigidly organized that the higher order arterial branches and arterioles were invariant (and they apparently have never been named). The gastric regions were, however, quite characteristic in terms of the general features of the major vessels.

The *antrum*, the muscular region of the distal stomach that grinds and macerates food mixed with gastric acid and enzymes and then pumps the resulting chyme into the duodenum, receives its primary supply from the right gastroepiploic artery. This artery, which courses on the greater curvature of the stomach, wraps small arterial branches around the body of the antrum. The higher order arterial branches were relatively small, apparently carrying a limited blood supply. A limited number of similarly small (or even smaller) arteries perfusing the antrum also course off the more rostrally located left and right gastric arteries associated with the corpus. Consistent with several estimates of the circulation required by muscle tissue (Kvietys, 2010, Peti-Peterdi et al., 1998), the antrum appears to receive small vessels presumably capable of supplying the circulation needed for its primarily muscular function.

By contrast, the *corpus* receives numerous larger arterial branches issued by the relatively symmetrically organized left and right gastric arteries. These large arteries wrap over the organ, coursing from the lesser to the greater curvature. As the large vessels course along the distal border of the corpus, numerous secondary arteries are issued as larger branches coursing rostrally and towards the greater curvature.

The secretory functions of the corpus mucosa require far more circulation than smooth muscle activity. (Peti-Peterdi et al., 1998), for example, found in an analysis of 25 rats that 84% of the blood flow to the stomach supported the secretory activity of the mucosa-submucosa (in the corpus region) whereas only relatively 16% of the blood flow supplied the smooth muscle of the stomach. Such dissimilar proportions are consistent with the additional fact that the corpus receives the blood flow of two paired and large gastric arteries, whereas the antrum is supplied primarily by only a single smaller vessel (the right gastroepiploic artery) and the forestomach similarly is perfused primarily by a single vessel (the left gastroepiploic artery).

Whereas the antrum is heavily muscular and the corpus is heavily secretory, the *forestomach* or fundus is specialized for storage of food until the nutrient material is moved into the corpus for the initial stages of digestion. The forestomach is less muscular than the antrum and (in the rat) is lined with squamous epithelium, not with mucosa and a dynamic epithelial layer such as is found in the corpus. In keeping with what must be less demand for circulatory exchanges, the forestomach is perfused by small arterial branches from the left gastroepiploic artery. In addition, the forestomach receives a few higher order arteriolar branches from the ipsilateral gastric artery.

### Factors Obscuring Arterial Patterns

In spite of this regular pattern, as illustrated in our Microfil defined observations (see Figures 1, 3 and 7), multiple factors in previous reports seem to have made the arterial pattern often harder to discern and should be noted. One factor that may well have contributed to some lack of attention to the vascular architecture, as well as use of overly schematic representations of the stomach vasculature, is blood pressure. Frequently, both experimental animal stomachs and human stomachs have been drawn or schematized using post mortem or cadaver material to estimate features. Since blood pressure is at zero, or near zero, in such cases, the larger vessels are likely to be partially collapsed and the finest vessels may even be completely drained of blood cells and plasma. In all likelihood, additional, accurate representations of the stomach vasculature will need to be done with fill agents or other means of generating normal pressures and contrasts in the vessels. In light of this consideration, it is relevant that we found that there was a window of pump circulation rates and pressures for Microfil infusions into the vasculature. Slower perfusion speeds and/or smaller volumes of contrast agent left the major vessels only modestly full and the finer arteries often unfilled; higher perfusion speeds and/or volumes often produced the formation of swollen blebs or “aneurysm-like” dilations, particularly in the finer vessels of the gastric mucosa, that appeared artifactual.

Furthermore, in the case of *in vivo* experiments, physiological factors operate to modulate vascular tone and often reduce vessel diameter to shunt blood away from some sites or to other regions both within the stomach and even between the different tissues and viscera. Physiological factors such as tissue oxygenation, hormones, sympathetic nervous system transmitters, paracrine signals released in the stomach wall, etc. can all change the vascular constriction or dilation pattern in a particular drainage field in the tissues cf., for example, (Kvietys, 2010). Such factors, which presumably normally work to modulate regional functions and link the food in the GI tract to the various need states and environmental stimuli the organism is negotiating at a given time. Certainly, factors influencing vasoconstriction and dilation may modulate the prominence of the stomach angioarchitecture during a surgical exposure as well as during normal physiological adjustments to local and immediate energy balance demands.

Finally another set of factors originating from the limited work on the blood vessels of the rat stomach, seem to have impeded analyses of the rat stomach angioarchitecture. The limited examinations of the rat and, in some discussions, the tendency to assume that the arterial patterns of rat and dog and human and other species are effectively interchangeable or can all be summarized with the same uncritical cross-species description may have confused the attempts to evaluate circulation specifically in the rat.

## Supporting information

Supplemental Video Animation 1

Supplemental Video Animation 2

## Acknowledgments and Funding Information

We thank Dr. Bartek Rajwa for extensive expert help with biostatistics and with modeling gastric emptying; Dr. Andy Schaber, Director of the Imaging Facility, Purdue’s Bindley Bioiscience Center for discussions of the microCT analyses; and Ms. Jennifer McAdams for all of her help with laboratory operations. As per NIH reporting guidelines, the research reported in this publication was supported by NIH DK27627 and by the Office of the Director, NIH under award OD023847.

## Author Contributions

T.L.P. and D.M.J conceived and designed research; L.C. performed experiments; D.M.J. analyzed data and interpreted results of experiments; D.M.J. and L.C. prepared figures; T.L.P. drafted manuscript; T.L.P. and D.M.J. edited and revised manuscript; T.L.P., D.M.J. and L.C. approved the final version of the manuscript.

## Compliance with Ethical Standards

No conflicts of interest, financial or otherwise, are held by any of authors. The present research involved animal subjects, and all animal husbandry practices conformed to the NIH *Guide for the Care and Use of Laboratory Animals* (8^th^ edition) and were reviewed and approved by the Purdue University Animal Care and Use Committee. All efforts were made to minimize any suffering as well as the number of animals used.

## SUPPLEMENTARY MATERIAL

**Supplementary Material 1** Video animation of a 3D image of the vasculature shown in Figure 1. The image was generated in 3D Slicer, and one full rotation (manually generated) of the image around the vertical axis (LES to GC) was captured using the screen recording tool of Camtasia®. The video was edited to loop after 20 seconds. DOI: 10.6084/m9.figshare.13970249

**Supplementary Material 2** Video animation of a 3D image of the segmented vasculature shown in Figure 2, upper panel. The image was created in Vesselucida® and one full rotation of the image around the axis of the LES to GC was recorded using the video creation tool in Vesselucida®. The video was edited to loop after 20 seconds. DOI: 10.6084/m9.figshare.13970528

**Supplementary Material 3** A table of model parameters and various assessments of model quality for four models of stomach emptying (two exponential versions and two logistic versions). Separate models are included for the forestomach and antrum-corpus compartments and for the total stomach. The second of the four models corresponds to the illustrative lines included in Figure 5. DOI: 10.6084/m9.figshare.13970789

## Notes

### Competing Interest Statement

The authors have declared no competing interest.

