## Supplemental Table (3) for "Stomach serosal arteries distinguish gastric regions of the rat"

The table below shows the parameters and measures of model fit for four different models for emptying of forestomach, antrum/corpus and total stomach. In the following equations v=compartment volume, M=meal size, W=animal weight and T=delay time from meal to perfusion, and a through e are model parameters.

The four models are:

#1: exponential decay model with meal size and delay time as variables

#2: exponential decay model with animal weight, meal size and delay time as variables

#3: logistic model with animal weight, meal size and delay time as variables

#4: logistic model with animal weight, meal size and delay time as variables, but with an added term in meal size.

The models are assessed by standard measures of model quality, except for Δ est-AIC which is quantified as the difference in 2k + n*ln(SSE) between models where n=number of data points and k=number of parameters excluding constants. There are, of course, many other models that could be evaluated, but quantitative analysis of stomach emptying is not the focus of this paper. However, the data below is included since these four relatively simple models were the strongest of those that were evaluated. Note also that the range of animal weight in the experiment was intentionally small, but, as the data below demonstrates, addition of weight as a variable does improve the quality of fit. Calculated emptying rates shown in the body of the paper were calculated using the exponential decay model with weight included to capture the empirical observations of emptying behavior without proceeding to the level of trying to define a realistic biophysical model of emptying.

| | | parameters | | RMSE | SMAPE | Δ est-AIC | Adj. Pseudo-R^2 |
|---|---|---|---|---|---|---|---|
| Forestomach | | | | | | | |
| #1 | v=c + b*M*exp(a*T) | a | -0.32 | 560.5 | 0.27 | 0 | 0.933 |
| | | b | 628 | | | | |
| | | c | 279 | | | | |
| #2 | v=a*W + b*M*exp(c*T + d*W) | a | 1.04 | 539.3 | 0.27 | -2.4 | 0.935 |
| | | b | 1127 | | | | |
| | | c | -0.32 | | | | |
| | | d | -0.0024 | | | | |
| #3 | v=c*W + (d*M – c*W) / (1+exp(b*W*(T-e))) | b | 0.0018 | 526.7 | 0.23 | -5.0 | 0.939 |
| | | c | 1.53 | | | | |
| | | d | 1781 | | | | |
| | | e | -1.41 | | | | |
| #4 | v=c*W + (d*M – c*W) / (1+exp(b*W*(T-e*M))) | b | 0.0022 | 536.5 | 0.21 | -3.0 | 0.937 |
| | | c | 1.80 | | | | |
| | | d | 1181 | | | | |
| | | e | 0.024 | | | | |
| Antrum-Corpus | | | | | | | |
| #1 | v=c + b*M*exp(a*T) | a | -0.23 | 583.2 | 0.12 | 0 | 0.981 |
| | | b | 376 | | | | |
| | | c | 2143 | | | | |
| #2 | v=a*W + b*M*exp(c*T + d*W) | a | 8.99 | 553.7 | 0.11 | -3.8 | 0.982 |
| | | b | 413 | | | | |
| | | c | -0.23 | | | | |
| | | d | -0.00040 | | | | |
| #3 | v=c*W + (d*M – c*W) / (1+exp(b*W*(T-e))) | b | 0.0017 | 547.5 | 0.11 | -5.1 | 0.982 |
| | | c | 9.91 | | | | |
| | | d | 929 | | | | |
| | | e | 0.22 | | | | |
| #4 | v=c*W + (d*M – c*W) / (1+exp(b*W*(T-e*M))) | b | 0.0017 | 547.6 | 0.11 | -5.1 | 0.982 |
| | | c | 9.89 | | | | |
| | | d | 958 | | | | |
| | | e | 0.0021 | | | | |
| Total Stomach | | | | | | | |
| #1 | v=c + b*M*exp(a*T) | a | -0.28 | 848.1 | 0.13 | 0 | 0.988 |
| | | b | 1000 | | | | |
| | | c | 2449 | | | | |
| #2 | v=a*W + b*M*exp(c*T + d*W) | a | 10.13 | 824.8 | 0.12 | -1.1 | 0.989 |
| | | b | 1519 | | | | |
| | | c | -0.28 | | | | |
| | | d | -0.0017 | | | | |
| #3 | v=c*W + (d*M – c*W) / (1+exp(b*W*(T-e))) | b | 0.0018 | 796.0 | 0.11 | -5.1 | 0.989 |
| | | c | 11.46 | | | | |
| | | d | 2444 | | | | |
| | | e | -0.48 | | | | |
| #4 | v=c*W + (d*M – c*W) / (1+exp(b*W*(T-e*M))) | b | 0.0020 | 794.7 | 0.10 | -5.3 | 0.989 |
| | | c | 11.75 | | | | |
| | | d | 2115 | | | | |
| | | e | 0.023 | | | | |
